# A general method to predict the effect of single amino acid substitutions on enzyme catalytic activity

**DOI:** 10.1101/236265

**Authors:** Yu-Hsiu T. Lin, Cheng Lai Victor Huang, Christina Ho, Max Shatsky, Jack F. Kirsch

**Affiliations:** Department of Molecular and Cell Biology, University of California, Berkeley, CA, USA; Department of Electrical Engineering and Computer Science, University of California, Berkeley, CA, USA; Physical Biosciences Division, Lawrence Berkeley National Laboratory, Berkeley, CA, USA; QB3 Institute, University of California, Berkeley, CA, USA

**Keywords:** Machine learning, bioinformatics, protein structure, site-directed mutagenesis, enzyme catalytic activity, enzyme active site

## Abstract

Over the past thirty years, site-directed mutagenesis has become established as one of the most powerful techniques to probe enzyme reaction mechanisms^1-3^. Substitutions of active site residues are most likely to yield significant perturbations in kinetic parameters, but there are many examples of profound changes in these values elicited by remote mutations^4-6^. Ortholog comparisons of extant sequences show that many mutations do not have profound influence on enzyme function. As the number of potential single natural amino acid substitutions that can be introduced in a protein of *N* amino acids in length by directed mutation is very large (19 * N), it would be useful to have a method to predict which amino acid substitutions are more likely to introduce significant changes in kinetic parameters in order to design meaningful probes into enzyme function. What is especially desirable is the identification of critical residues that do not contact the substrate directly, and may be remote from the active site.

We collected literature data reflecting the effects of 2,804 mutations on kinetic properties for 12 enzymes. These data along with characteristic predictors were used in a machine-learning scheme to train a classifier to predict the effect of mutation. Use of this algorithm allows one to predict with a 2.5-fold increase in precision, if a given mutation, made anywhere in the enzyme, will cause a decrease in k_cat_/K_m_ value of ≥ 95%. The improved precision allows the experimentalist to reduce the number of mutations necessary to probe the enzyme reaction mechanism.

## INTRODUCTION

Site-directed mutagenesis has emerged as the most precise probe of the role of individual amino acid side chains in the catalytic mechanism of an enzyme^1–3,7–10^. Normally, a target residue is replaced with one of the other 19 natural amino acids, and the catalytic activity of the mutant is compared with that of the wild-type (WT) enzyme. Interpretation of the experimental result may be complicated. Interestingly, a negative result is usually definitive in that it allows the conclusion that the original side chain residue has no important catalytic function. The interpretation of a positive result, wherein the mutation leads to a significant loss in activity, to imply that the probed side chain is intimately involved in the catalytic machinery may be compromised. It is possible that the original side chain, for example, helps to maintain the overall architecture of the active site. Replacements of a target amino acid with several others, rather than with a single one, often yield variants with different activity levels^11^. A notable example is seen in probes of the paradigmatic Asp-His-Ser catalytic triad of the serine protease, subtilisin. Replacement of Asp-32 with Ala produced a variant that is 10^-4^ times as active as the WT^12^. This was taken as further support for the existence of a short strong H-bond between Asp-32 and His-64 and for its importance in catalysis. However, subsequent substitution of Asp-32 with Cys removed the short strong H-bond, but this construct retained 12% of the WT activity in terms of k_cat_/K_m_^9^.

Nearly all rationally introduced mutations, which are designed to elucidate mechanism, are deliberately introduced at the active site, and indeed substitutions of those side chains intimately involved in catalytic mechanism or in substrate binding nearly invariably do result in severe compromise of catalytic activity. However in the absence of prior knowledge it would be very difficult to identify positions remote from the active site that are likely to result in significant decrements in the catalytic parameters. Nonetheless several examples of remote introduced mutations that do have significant effects on catalytic activity have been obtained^5,6,13^, and these often provide important insights into, for example, the coupling of dynamics with catalytic activity ^14,15^.

A complete experimental exploration of all possible single amino acid substitutions in a given protein requires considerable experimental effort and is currently expensive; therefore, it would be useful to have a method to predict which amino acid substitutions are more likely to introduce significant changes in kinetic parameters in order to design meaningful probes into enzyme function. What is especially desirable is the identification of critical residues that do not contact the substrate directly, and may be remote from the active site. A computational procedure that would significantly increase the probability of correctly predicting, inter alia, severely deleterious remote mutations, would correspondingly reduce the experimental effort required for such identification. We collected data reflecting effects of 2,804 mutations on kinetic properties for 12 enzymes. These mutation data along with characteristic predictors, which define each substitution, were organized in a MySQL database. The predictors include distance from the active site, position-specific amino acid frequency scores (PSAAFS) on related sequences, and natural substitution frequency (BLOSUM62)^16^. The predictive power for the effect of a novel mutation was evaluated independently for each of the three predictors, for the three possible combinations of any two of them, and for all three together. The last classifier, incorporating all three predictors performed significantly better. Use of this algorithm allows one to predict with a 2.5-fold increase in precision, whether a given mutation, made anywhere in the enzyme, will affect a decrease in k_cat_/K_m_ value of ≥ 95%, thus permitting significant reduction in experimental effort necessary to identify deleterious mutations.

## MATERIALS AND METHODS

### Data collection

#### Literature search

A large database representing the effects of single amino acid substitutions on enzyme activity was assembled to serve as a training set for classifier applications. The Protein Mutant Database^17^, PubMed^18^, and Google Scholars^19^ were used to gather mutations on enzymes that satisfy the following criteria: 1) the availability of at least one structure complexed with a substrate or inhibitor in the PDB and 2) lack of reported allosteric behavior. The effects of 2804 mutations on the kinetic properties of 12 enzymes were acquired. **Table 1** lists the enzymes and associated mutations included in the MySQL database. Data are available for at least 14 mutations from each of eight enzymes. The collected data include the following information: the substituted mutation, mutant kinetics (k_cat_/K_m_), wild-type kinetics (k_cat_/K_m_), organism, and substrate. The structure of the database is depicted in **Figure S1**.

**Table 1.** Enzymes included in the analysis, sorted by number of mutations*.

| Enzyme | Sequence length | # of mutations | Substrates | PDB ID | Refs. |
| --- | --- | --- | --- | --- | --- |
| T4 lysozyme (T4L) | 164 | 1977 | *N*-acetylglucosamine, *N*-acetylmuramic acid, peptide | 148L | 5 |
| HIV protease | 99** | 478 | Dihydroethylene-containing inhibitor | 4FL8 | 6 |
| Thymidylate synthase (TS) | 316 | 173 | Deoxyuridine monophosphate (dUMP), tetrahydrofolate (THF) | 1LCA | 11,35–44 |
| Dihydrofolate reductase | 159 | 50 | 7, 8-dihydrofolate, NADPH | 3DGA (*P. falciparum*), 1RX3 (*E. coli*) | 4,13,45–49 |
| Aspartate aminotransferase | 396** | 44 | 2-oxoglutarate, L-Asp, L-Glu, oxalacetate | 1ASM | 50–60 |
| Glutathione transferase | 219 | 34 | 1-chloro-2,4-dinitrobenzene, L-glutathione | 1JLW | 61 |
| Salutaridine reductase | 311 | 22 | NADPH, salutaridine | 3O26 | 62 |
| Hexokinase | 455 | 14 | ATP, glucose | 1V4S | 63 |
| Arginase | 323** | 5 | L-Arginine | 1RLA (*R. norvegicus*), 2AEB (*H. sapiens*) | 64 |
| Fatty acid amide hydrolase | 537** | 4 | Oleamide, methyl oleate | 1MT5 | 65 |
| Cytosine deaminase | 161** | 2 | 5-fluorocytosine, cytosine | 1P6O | 66 |
| Fructose-bisphosphate aldolase | 358** | 1 | Fructose 1,6-bisphosphate | 1DOS | 67 |

\*A total of 2804 mutations were collected for the 12 listed enzymes for analysis. At least 14 mutations are recorded for eight of the enzymes. \*\*For each monomer of the homodimer, or trimer as in the case of arginase.

### Predictors

Several predictors that were anticipated to correlate with loss of catalytic activity were computed for each substitution, and they are described below.

#### Distance from Active Site

The distances (Å) of each mutation from the enzymatic active site were calculated from the PDB files. They were measured from the catalytic site to the closest atom on the side chain of the residue of interest. The active site for each enzyme was determined from knowledge of the catalytic mechanism and the ligand binding coordinates. The method used for each enzyme and the justification are given in the **Table S1**.

#### Position-Specific Amino Acid Frequency Score (PSAAFS)

A multiple sequence alignment (MSA) of closely related protein sequences to an enzyme of interest can reveal the degree of conservation at each position and is thus informative for predicting the effect of an amino acid substitution on catalytic activity. A protein sequence along with a set of default parameters (database UniRef90 2011 April, median conservation of sequences: 3.00, remove sequences more than 90% identical) is first submitted to SIFT^20^, a sequence-based method developed to predict the effect of amino acid substitutions on protein function. The output MSA is then used to derive a set of position-specific amino acid frequency scores (PSAAFS) that fill an *m*-by-*n* matrix where *m* corresponds to the 20 canonical amino acids and *n* represents the length of the protein. The PSAAFS for residue *R* at position *n*, or *R_n_*, is computed using the following equation, where 1) *F_R,n_* is the ratio of the total number of times residue *R* occurs at position *n* to the number of aligned sequences, 2) *A_n_* is the number of unique amino acids present at position *n* in the alignment, and 3) *G_n_* represents the number of gaps at position *n* divided by the number of aligned sequences:

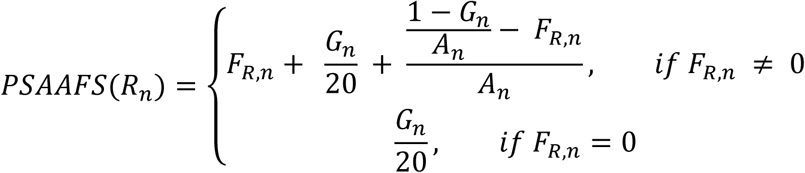

The formula above accounts for gaps in the alignment as well as number of unique amino acids present at each position. The first term, *F_R,n_*, denotes the proportion of times that residue *R* appears at position n. The second term evenly distributes the space taken up by the gaps at position *n* among the 20 residues. The final term reflects the number of distinct residues found at position n, or a rough consideration of entropy. For example, consider two cases where we would like to find the PSAAFS score for alanine. In an alignment of ten sequences at one position there is one alanine and nine leucines, while in the same alignment at another position there is one alanine, four leucines, and five valines. Even though alanine appears the same number of times in both cases, in the former instance we would like to assign a higher score for alanine because fewer unique residues are present at that position, and thus we consider that position to be more conserved. Without the third term in the equation, the score for alanine would be exactly the same, while PSAAFS assigns scores of 0.3 and 0.178, correspondingly. Altogether, the PSAAFS scores for all residues at a given position sums up to 1.

#### BLOSUM

BLOSUM^16^ is a matrix derived from the conservation of amino acid substitutions. Scores from BLOSUM62, which is based on an alignment of sequences with less than 62% sequence identity, were used. The scores represent the likelihood of a given substitution to occur naturally.

#### Other Predictors

3D-1D substitution matrix^21^, SIFT^20^, solvent accessibility^22^, sequences conservation scores computed from Pfam multiple sequence alignments^23^, and secondary structures^24^ were considered additionally, but their inclusion did not improve the prediction success.

### Classification

We explored various classifiers, such as J48, Random Forest and RandomTree, in the WEKA package^25^, and selected Random Forest, which consistently gave good performance. Random Forest is an ensemble classifier that constructs multiple decision trees, each of which results in a classification “vote.” These trees are constructed by: 1) bootstrapping from training set data and 2) selecting, at each node, the best split among a random subset of predictors. Aggregating the votes from these decision trees dictates the prediction outcome. This algorithm has proven to be robust in detecting intrinsic patterns between the provided predictors and some output decision^26^. In the present case, features of amino acid substitutions, such as distance from active site and conservation scores, are used to make a binary choice of whether the mutant retains more than 5% of its wild-type enzymatic activity. The results presented in this paper were computed using the “randomForest” package^27^ implemented in R^28^.

### Assessment of classification accuracy

The performance was measured using Receiver Operating Characteristic, or ROC curves, as well as precision plots. The data-set was classified as follows:

Positives: deleterious mutations (≥ 95% loss of activity).

Negatives: neutral mutations (< 95% loss of activity).

True Positives (TP): correctly predicted deleterious mutations.

False Positives (FP): neutral mutations predicted as deleterious.

ROC curves are one of the most commonly used ways to visualize accuracy of a classification procedure. The y-axis defines the true positive rate (TPR), which is the number of correctly predicted deleterious mutations over all true deleterious mutations. The x-axis defines the false positive rate (FPR), which is the number of neutral mutations incorrectly predicted as deleterious divided by all true neutral mutations. A curve rising steeply from the origin to the top-left corner represents a successful classification, while a random prediction corresponds to a diagonal running from the bottom-left to the top-right corner.

Additionally, precision plots were calculated to facilitate experimental design as they inform the experimentalist about the expected yield of deleterious mutations for any arbitrary number of total mutations introduced by site-directed mutagenesis. The y-axis is known as “precision”, i.e. TP/(TP+FP) while the x-axis is the number of deleterious predictions, i.e. TP+FP.

### Method validation

#### 10-Fold Cross-Validation (10FCV)

Cross-validation is a statistical method to determine the practical value of prediction models. It is a three-step process: 1) The dataset is divided into a training set and test set; 2) the model is fitted on the training set before making predictions on the test set; 3) the performance is evaluated by comparing the predictions to the experimental result. The dataset is divided into 10 subsets in 10-fold cross-validation (10FCV). Each subset is withheld in turn from training the model for testing and evaluation. This procedure is iterated 10 times, once for each subset.

A caveat with 10FCV involves the selection of subsets for cross-validation. Subsets should be determined such that the elements in each subset are independent to all other elements in the other subsets. For example, at position 12 on T4 lysozyme, substitutions to 12 different residues were characterized. If the subsets were randomly selected, it is likely that these 12 mutations will be found in different subsets. This creates a problem during cross-validation because different mutations made at the same position in a given enzyme frequently may have similar effects on catalytic activity. When training and testing sets contain related data points, the model would appear to perform better than it does in reality. The 10FCV applied on our mutation data includes an additional step to avoid this pitfall. All substitutions made on the same residue in the same enzyme are treated as a single element. These elements are then randomly divided into 10 subsets for 10FCV.

#### Leave-One-Enzyme-Out (LOEO)

In addition to 10FCV, a variation of cross-validation called leave-one-enzyme-out (LOEO) was employed for model assessment. Here the mutations were partitioned into subsets based on the probed enzyme. Next, all mutations associated with one enzyme were set aside for testing while the remaining data were used to train the classifier. The model then predicted the activity changes induced by mutations in the test set. We iterated through this process until all enzymes had been withheld for testing.

## RESULTS AND DISCUSSION

### Predictors of mutation-induced decrement in catalytic function

Several computational predictors were evaluated in order to estimate their utility to contribute to the determination of whether the effect of an arbitrarily chosen mutation is severely detrimental to the catalytic activity of a selected enzyme. After considerable experimentation, it emerged that the most useful predictors were distance from the active site (*r*), likelihood of pairwise substitutions (BLOSUM62)^16^, and position-specific amino acid frequency score (PSAAFS)^20^. We also evaluated 3D-1D scores from environmental profiles^29^, Pfam conservation scores^23^, solvent accessibility of residues^22^, and types of secondary structure^24^ as additional possible predictors. The latter evaluators added very little to what was obtained using only *r*, BLOSUM, and PSAAFS, and were not considered further.

### Benchmark data set

The objective of the present research is to develop a computation procedure to predict mutations in any enzyme with known structure and an identifiable active site that have a high probability of decreasing the k_cat_/K_m_ value by ≥ 95%. In order to evaluate the predictors individually and in combination we constructed a benchmark data set of known mutation induced changes in enzyme catalytic activity. We searched the literature for quantitative reports of changes in kcat/K_m_ resulting from mutation from the wild-type. In some cases, where k_cat_/K_m_ values were not available, we collected phenotypic descriptors and converted them into numeric values. The developed algorithm requires that an enzyme structure in complex with at least one known substrate or analog be available in the PDB^30^. **Table 1** lists the collected set of enzymes and the number of literature reported mutations for each. Historically the vast majority of introduced mutations that resulted in large decrements in k_cat_/K_m_ values were made at residues near the active sites in order to explore mechanistic hypotheses, and thus do not allow extensive exploration of the distance parameter. Fortunately, however, for the present exercise, every position in T4 lysozyme (T4L)^5^ and in HIV protease^6^ had been substituted with one or more residues, thus permitting a thorough examination of the *r* parameter in isolation and together with BLOSUM and PSAAFS. The database is thus dominated by the 1,977 and 478 mutations reported for T4L and for HIV protease, respectively. These two enzymes together comprise 88% of all of the collected mutations. The advantage of such an abundance of data from these two enzymes is, in part, offset by the fact that the effects on kinetic parameters were not investigated. However, Hardy et al.^5^ noted that viral viability requires at least 3% activity in T4L. We rounded this figure to the 95% decrease in activity discussed above. Swanstrom et al.^6^ scored HIV viability for the mutant proteases as positive, intermediate, and negative. Our initial assumption is that the negative scores also correspond to < 5% activity. The remaining enzymes reporting on the effect of mutations exhibit a wide range of catalytic activities. This enzyme collection was restricted to those with reported k_cat_/K_m_ values for small molecule substrates. The number of mutations per enzyme varies from 173 on thymidylate synthase (TS) to one on fructose-bisphosphate aldolase, as shown in **Table 1**.

### Method evaluation with ROC and precision plots

In order to estimate the effect of a given mutation on catalytic activity we applied a machine learning approach wherein a classifier was trained on the collected mutation data in our database using values of three predictors r, BLOSUM, and PSAAFS. We explored various classifiers in the WEKA package^25^ and selected Random Forest, which consistently gave good performance that was evaluated by receiver operating characteristic (ROC) plots **(Figure 1)**. These plots depict the accuracy of the predictions by relating the true positive rate (TPR) to the false positive rate (FPR). TPR is defined as the ratio of true positives to the sum of true positives and false negatives TP / (TP + FN); FPR represents the ratio of false positives to the sum of true negatives and false positives FP / (TN + FP). Here, positives are defined as mutations that result in at least 95% loss of catalytic activity, while negatives are categorized as neutral mutations that retain >5% of the wild-type function. A classification able to separate positives and negatives perfectly would result in a step-shaped ROC curve that initiates at the origin, rises vertically to the coordinate (1,0), and terminates with a horizontal line to coordinate (1,1). The area under such a curve (AUC) equals one. Conversely, randomly generated classifications with no predictive value will exhibit a diagonal line connecting (0,0) to (1,1) with an AUC of 0.5^31^. **Figure 1a** shows a ROC plot constructed by using the reported data for T4L as a training set to predict deleterious mutations in HIV protease. The top curve (solid line) delineates an AUC of 0.806, which is considerably better than random. To illustrate - an experimentalist willing to accept an FPR of only 0.1, would expect a TPR of 0.6, which is six-fold better than a random prediction that would yield a TPR of 0.1.

**Figure 1.**
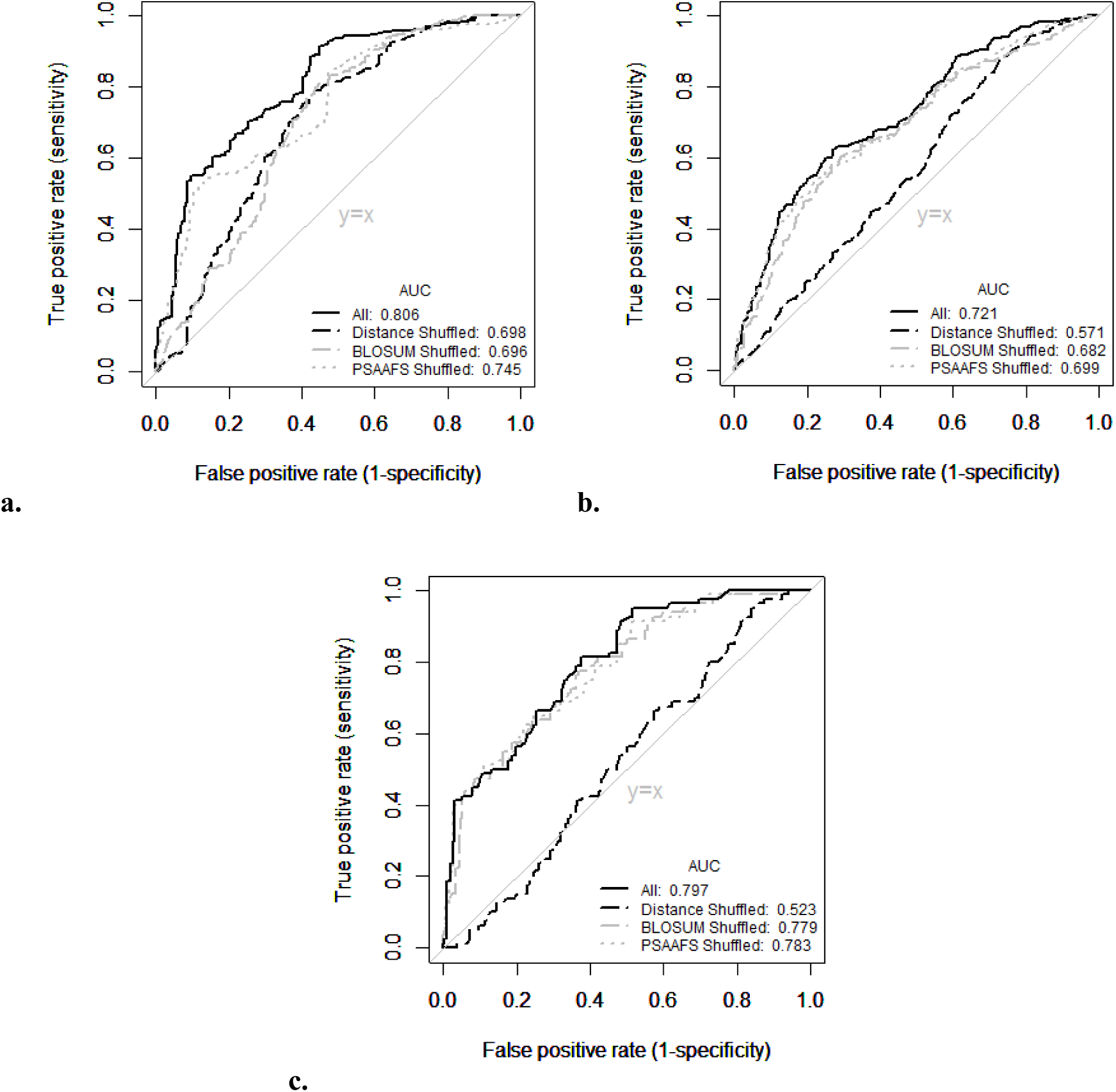
Estimated false positive and true positive rates as a function of shuffling each predictor individually. The performance is measured from **(a)** training on T4L mutant data set to test on HIV protease, **(b)** training on HIV protease mutants to predict T4L mutation results, and **(c)** training on T4L and HIV protease data sets combined to predict effects of mutation on all 10 other enzymes. In the legends, “All” indicates that all three predictors, distance, BLOSUM, and PSAAFS, were used in training and prediction. The others were constructed by shuffling the indicated predictor in order to assess its contribution. The areas under the curves (AUC) are given in the insets.

The curves in **Figure 1** illuminate that the greater contribution of distance to the overall performance compared to the other predictors. **Figures 1b and c**, respectively, were constructed from training on HIV protease data to predict T4L mutations, and training on the combined T4L and HIV protease data sets to predict the effects of mutations on the remaining ten enzymes. For each case, the individual parameters were shuffled in turn to evaluate their predictive abilities. In

**Figure 1c**, the AUC decreased from 0.797 to 0.523, when distance was shuffled as a predictor, while much smaller decrements resulted from scrambling either BLOSUM or PSAAFS data. The same observation was made from **Figures 1b and c** to demonstrate the impact of distance on the method performance.

Although ROC curves provide a good estimate of the performance of prediction algorithms, they may mislead, particularly where the number of negative outcomes (i.e. neutral mutations) is significantly higher than the number of positive ones (i.e. deleterious mutations)^32^. An illustrative example is taken from **Figure 1b**, which summarizes the method performance from training on HIV protease mutation data to predict the effect of mutations on T4L activity. The experiments on mutagenesis on T4L detected 280 (14.2%) deleterious and 1697 (85.8%) neutral mutations. An AUC of 0.721 suggests that this procedure produces substantially more accurate predictions than would be expected by chance; however, a deeper examination reveals a poor performance when predicting deleterious mutations. For example, if an experimentalist were to allow an FPR of 0.05, it means that only ~85 (5%) of the 1697 neutral mutations, would be predicted incorrectly as deleterious. The corresponding TPR of 0.20 indicates that only 56 (20%) would be predicted correctly as deleterious. **Table 2** presents an error matrix that provides a breakdown of the model performance at an FPR threshold of 0.05.

**Table 2.**
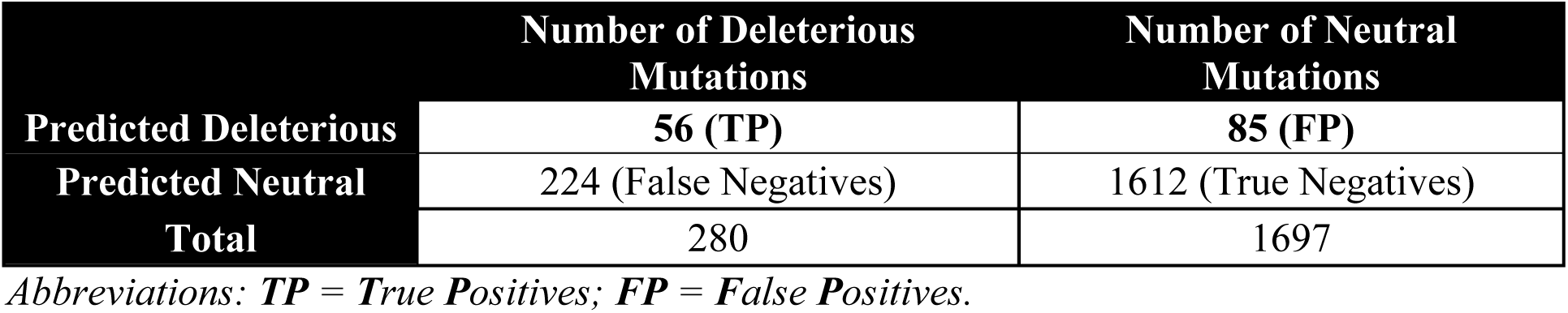
Error matrix of predictions on T4L mutants at 0.05 FPR.

Thus a small FPR, which is usually interpreted as a highly accurate classification, in reality may result in low precision, which is defined as TP / (TP + FP). Precision represents the fraction of predicted deleterious mutations that is correctly classified. The above example shows that, out of the total 141 (56 + 85) mutations predicted to be deleterious, 56 were scored correctly, resulting in a precision of only 40% despite having selected a stringent FPR threshold. In other words, while a method may incorrectly classify a small percentage of neutral mutations as deleterious (here 85 / (85 + 1612)), their absolute number might be so large that the overall number of predicted deleterious mutations may be dominated by false positives.

Precision plots of TP / (TP + FP) vs. TP + FP supplement ROC curves and are especially useful when one class, here the negatives (neutral mutations), is significantly overrepresented. Precision plots offer an additional advantage in that they may provide better guidance for experimentalists, who are usually interested in maximizing the yield of deleterious mutations for a given number of constructed mutants as perturbing mutations are more likely to inform on the mechanism of action of the probed enzyme. Precision plots can guide the experimentalist in choosing to accept a certain number of top predictions as deleterious mutations. For example, **Figure 2a** shows such a plot of predictions on T4L after training on the mutational data for all 11 of the other enzymes. Of the top 100 predicted deleterious mutations from the Random Forest classifier (See Methods), 40 would be correctly identified as contributing to losses of function. The precision plot also shows a “random precision” line that denotes the fraction of predicted deleterious mutations correctly assigned when predictions are made at random, where only 14 of the top 100 mutations would have been correctly identified.

**Figure 2.**
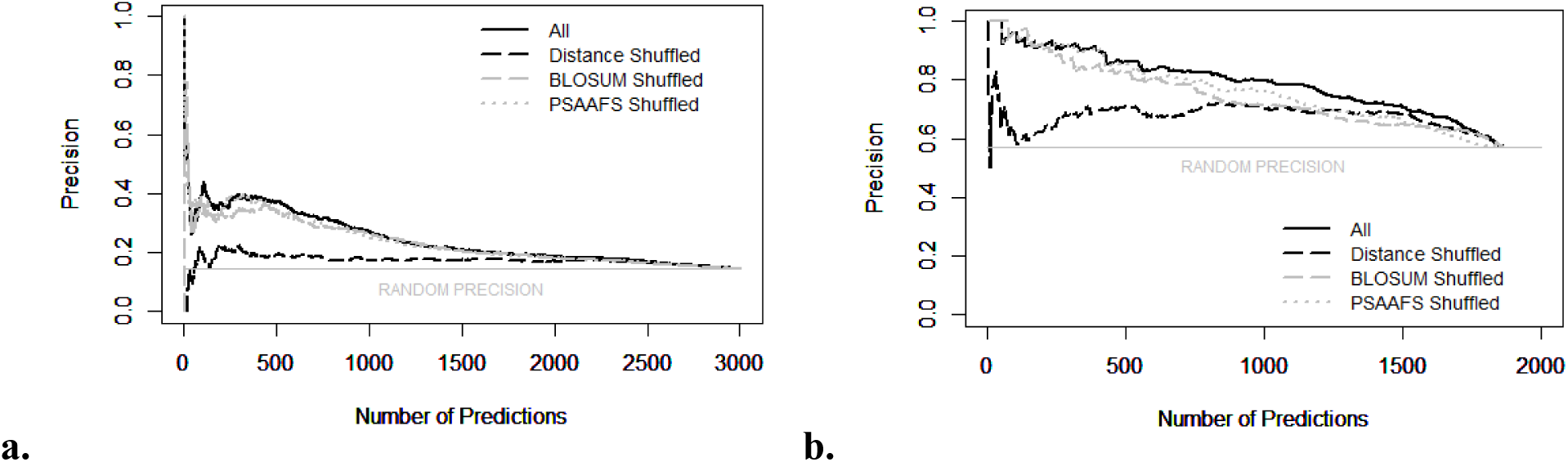
Precision curves as a function of shuffling each predictor individually. Predictions were made for all possible mutations on either **(a)** T4L or **(b)** HIV protease. Each example was trained on the mutation data for all 11 other enzymes. The predicted mutations were scored by the Random Forest classifier and ranked in order of likelihood to be deleterious. Each predictor was independently shuffled to assess its overall contribution to the method performance, as was done for the ROC curves. The “Random Precision” line shows the fraction of experimentally reported deleterious mutations.

The utility of a given precision plot or ROC curve is, of course, a function of the data used in the training set and of the choice of target enzyme. For example, **Figure 2b** shows a precision plot predicting deleterious mutations in HIV protease using the data for the 11 other enzymes in our data set for training. The performance here is much better than that shown in **Figure 2a**. A precision of 0.962 for the 100 highest-scoring HIV protease mutants was achieved compared to a precision of 0.4 computed for the 100 top-scoring T4L mutants. The difference in outcomes between **Figures 2a and 2b** is likely a result of the fact that 57.1% of the reported mutations in HIV protease are deleterious while only 14.2% of those reported for T4L are.

ROC curves and precision plots provide two distinct ways for model evaluation, and both demonstrate that the distance parameter has the greatest contribution to the overall performance of our method. In line with our findings with ROC curves, the precision plots in **Figure 2** clearly show a greater decrease in performance when the distance predictor is shuffled compared to shuffling BLOSUM or PSAAFS. This suggests that the distance parameter is critically important for accurate predictions.

### Evaluation of method performance using cross-validation (CV)

An ideal evaluation procedure would be to select an enzyme for which no mutagenesis data on catalytic activity are available, apply the above-described procedure, and test the predictions experimentally. Alternatively established methods can be applied to the existing data sets to achieve a similar goal. One of which is the 10-fold cross-validation^33^ (10FCV). In this procedure, the data are first randomly partitioned into 10 subsets; then each subset is withheld in turn for testing while the other nine are reserved for training. The 10FCV approach assumes that the data points are independent samples from the underlying distribution. However, in the present case, the mutation data are not entirely independent because often several substitutions were introduced at the same position. Therefore, 10FCVs have to be exercised judiciously. **Figure 3** presents two sets of ROC curves, one of which was executed on the **(a)** T4L mutant data and the other on the **(b)** HIV protease data set. The ROC curves for 10FCV in **Figure 3** were constructed by stratifying the data such that no single residue appears in more than one subset even if more than one mutation had been evaluated at the position. The second procedure that was employed to test the performance of a method is Leave-One-Enzyme-Out cross validation (LOEO), where each of the enzymes was independently excluded from the training set for evaluation. This approach is represented alongside 10FCV in **Figure 3** on the **(a)** T4L and **(b)** HIV protease data sets. As shown, 10FCV performs equally to LOEO for both T4L and HIV protease in predicting the effects of mutations. The AUC for T4L in 10FCV (0.702) is a little less than in LOEO (0.721). Likewise, the AUC for HIV protease in 10FCV (0.783) is only slightly lower than in LOEO (0.797). The similarity in the outcome of the two evaluation approaches validates the stability of the classification.

**Figure 3.**
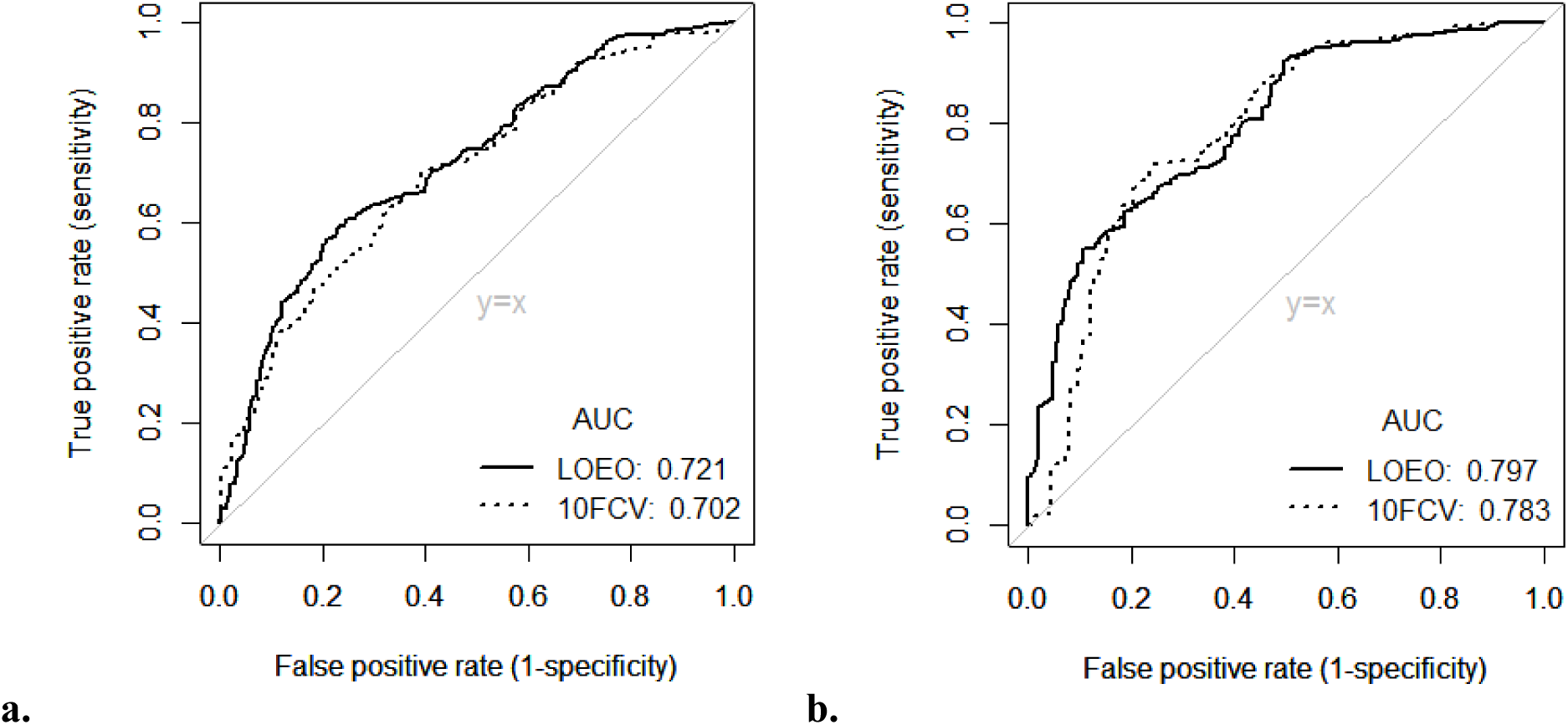
ROC curves comparing Leave-One-Enzyme-Out (LOEO) and 10-fold cross-validation (10FCV). These evaluation procedures were applied on the **(a)** T4L and **(b)** HIV protease mutant data sets. The distance, BLOSUM62, and PSAAFS predictors were used in both cross-validation analyses. See text for details.

To provide a more comprehensive assessment of the method, the three enzymes with the largest number of mutants in the database—T4L, HIV protease, and TS—were selected for LOEO evaluation. **Figure 4** presents precision plots from LOEO for predicting the effects of mutations on **(a)** T4L, **(b)** HIV protease, **(c)** TS with substrate dUMP, and **(d)** TS with substrate THF. As expected, the success rate is higher for enzymes having a greater percentage of deleterious mutations. Thus, the method performance was evaluated through a comparison of the precision plot to the baseline, represented by a solid gray line on each plot. The 100 top-scoring predictions from Random Forest for deleterious mutations for each enzyme were selected for further analysis. A precision of 0.4 over the baseline of 0.142 was realized for T4L mutants, which is a three-fold improvement over a random outcome. For predictions on the HIV protease, a precision of 0.962, well above the baseline of 0.571, was achieved. Precisions of 0.938 and 0. 760 were attained over random precisions of 0.734 and 0.603 with substrate dUMP and THF, respectively, for predictions on TS mutations. While the method improves the prediction of deleterious mutations for the enzymes presented above, the variability in these improvements over their respective baselines is dictated by both the complexity of the task and the ratio of deleterious mutations to neutral ones. The statistical significance of our predictions versus random is revealed through the non-parametric test of ROC curves by Delong^34^, which gave P-values of 1.11e-7, 7.31e-23, 5.44e-4, and 7.62e-5 that support a statistically significant improvement of predictions on T4L, HIV protease, TS with dUMP, and TS with THF, respectively, over random estimations.

**Figure 4.**
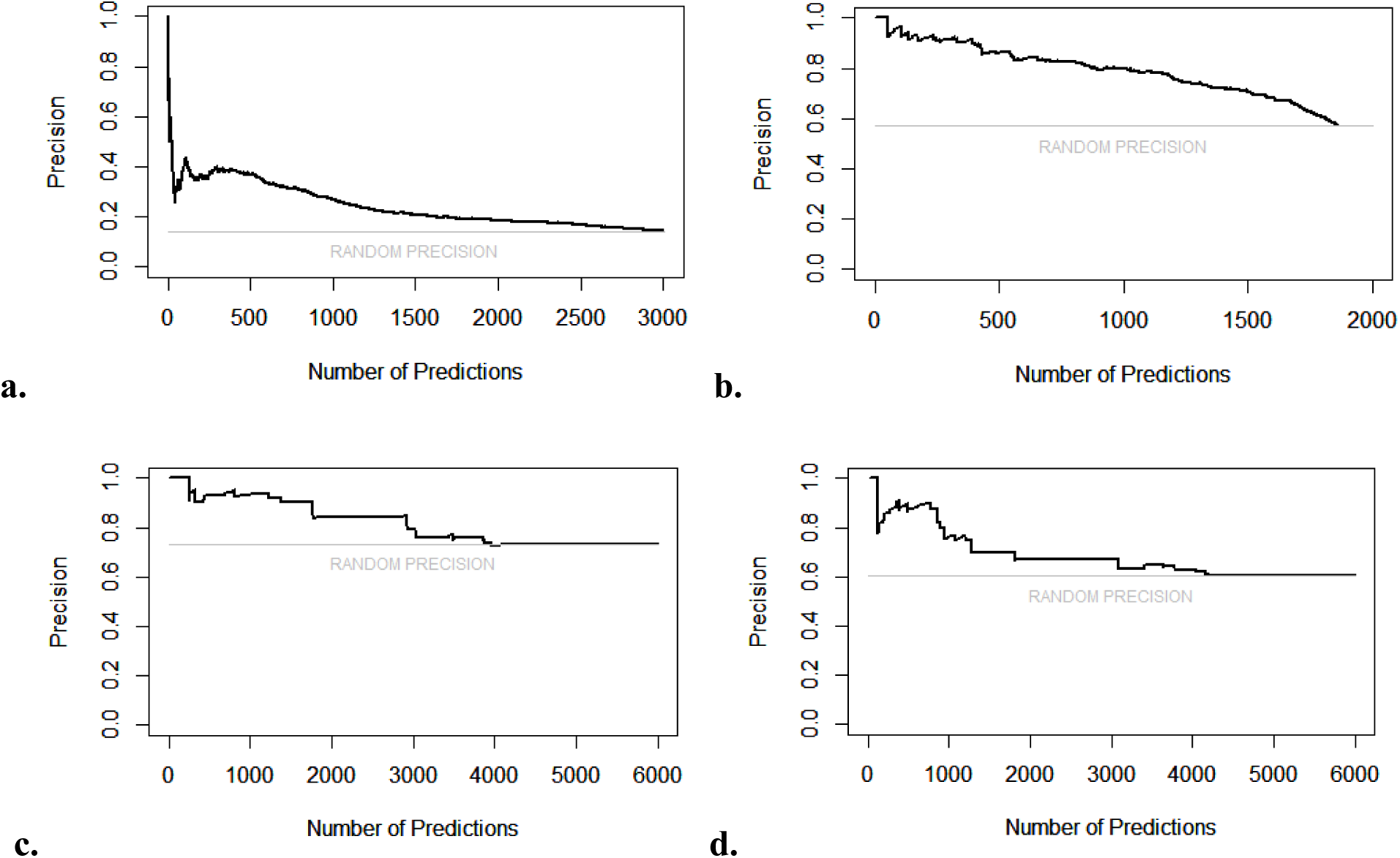
Precision plots calculated for Leave-One-Enzyme-Out cross-validation (LOEO) on specific enzymes. The evaluation was performed on the three enzymes for which the most data are available—**(a)** T4L, **(b)** HIV protease, and TS with **(c)** dUMP as the substrate and **(d)** with THF. The training sets contained all mutation data from **Table 1** excluding those for the tested enzyme. The distance, BLOSUM62, and PSAAFS predictors were used in generating the model. Precision curves were constructed for all eight enzymes where data for at least 10 mutations were available. Curves for the remaining five are not shown to maintain clarity.

### Sample predictions from Random Forest

An experimentalist wishing to illuminate the most kinetically damaging mutations in an enzyme, would need to provide the protein sequence, the PDB structure, and the active site coordinates as determined from mechanistic analysis. The algorithm employs Random Forest to calculate the probability of a ≥ 95% decrease in k_cat_/K_m_ for all possible natural mutations.

Random Forest predictions range from 0 (likely neutral) to 1 (likely deleterious). For example, a score of 0.70 for a given mutation indicates that 70% of the *n* (1000 used here) trees constructed by the ensemble classifier, call the mutation deleterious; therefore the remaining (30%) of the trees classify it as neutral. An output near 0 or 1 implies increased certainty that the mutation results in no change or a loss in function, respectively. **Table 3** shows a sample Random Forest output from applying LOEO individually for the four enzymes with the greatest numbers of mutations. The training set contained all mutations from the 11 enzymes that had not been omitted. For demonstration, we set an arbitrary cutoff at 0.5 to classify mutations above or below that score as deleterious or neutral, respectively. The table displays 20 mutations for each enzyme in descending order of certainty, starting with the most confident deleterious prediction and extending to neutral predictions for only TS and AspAT. Since the database contains a large number of T4L and HIV protease mutations, predictions for only the 20 top-scoring deleterious mutations are displayed for these two enzymes. The experimental results (ER) column denotes the experimental evidence for the mutations to be classified as **N**eutral or **D**eleterious. The Random Forest predictions are listed under the **Pred** column. For example, the first listed mutation for T4L, Q105C, was predicted correctly to be deleterious with support from more than 91% of the constructed trees. Conversely, Q105G was predicted incorrectly with 90.7% of the classification trees scoring it as deleterious when it is in fact neutral. Comparing the results across the four enzymes, we saw, as expected, that the quality of the prediction is higher when an enzyme with few mutations is omitted from the training set, e.g. AspAT, compared to when an enzyme carrying a large number of mutations is excluded, e.g. T4L.

**Table 3.** Sample Random Forest predictions for T4L, HIV protease, TS (with dUMP and THF), and AspAT using the LOEO procedure. Distance, BLOSUM62, and PSAAFS were used as predictors. The experimental results (ER) column shows the experimental evidence for the mutations as either deleterious (D) or neutral (N). The table is ordered by the Prediction (Pred) column from high (likely deleterious) to low (likely neutral). All predictions for deleterious mutations (Pred > 0.50) are in bold.

| T4L | ER | Pred | HIV | ER | Pred | TS_dUMP | ER | Pred | TS_THF | ER | Pred | AspAT | ER | Pred |
| --- | --- | --- | --- | --- | --- | --- | --- | --- | --- | --- | --- | --- | --- | --- |
| Q105C | D | **0.912** | A28S | D | **0.76** | H199V | D | **0.619** | R23F | D | **0.547** | W140F | D | **0.756** |
| Q105F | D | **0.912** | G27A | D | **0.735** | H199A | D | **0.608** | R23I | D | **0.547** | W140G | D | **0.721** |
| R145C | N | **0.912** | G27I | D | **0.724** | H199T | D | **0.608** | R23V | D | **0.547** | Y70S | D | **0.721** |
| R145F | D | **0.912** | G27L | D | **0.724** | R23F | D | **0.544** | W82A | D | **0.508** | Y225R | D | **0.718** |
| Q105G | N | **0.907** | D25H | D | **0.706** | R23I | D | **0.544** | W82S | D | **0.508** | A224I | N | **0.715** |
| Q105L | N | **0.907** | D25Y | D | **0.704** | R23V | D | **0.544** | W82N | D | **0.508** | K258H | D | **0.709** |
| R145G | D | **0.907** | G27V | D | **0.704** | H199S | D | **0.511** | E60L | D | 0.487 | K258A | D | **0.704** |
| R145L | D | **0.907** | G27D | D | **0.703** | R218K | D | **0.504** | P197L | D | 0.480 | H143A | N | 0.48 |
| R145Y | D | **0.907** | L23P | D | **0.699** | E60L | D | 0.484 | P197Y | D | 0.480 | R292D | D | 0.467 |
| R145P | N | **0.907** | D25A | D | **0.695** | D221C | D | 0.482 | P197C | N | 0.480 | C191A | N | 0.437 |
| T142G | N | **0.905** | G27P | D | **0.695** | P197I | D | 0.475 | P197I | N | 0.480 | C191S | N | 0.397 |
| T142Y | D | **0.905** | G27R | D | **0.695** | P197L | D | 0.475 | H199V | D | 0.473 | D222E | D | 0.392 |
| T142F | N | **0.905** | L23V | N | **0.686** | P197Y | D | 0.475 | W82F | D | 0.469 | Y70F | D | 0.345 |
| T142H | D | **0.905** | A28G | D | **0.61** | P197C | N | 0.475 | R23Q | D | 0.467 | Y225F | N | 0.345 |
| R14C | N | **0.905** | A28T | D | **0.61** | P196C | D | 0.472 | P197F | D | 0.449 | V39L | N | 0.296 |
| R14F | N | **0.905** | L23Q | D | **0.607** | P196I | D | 0.472 | P197W | D | 0.449 | C192A | N | 0.296 |
| Y18G | D | **0.902** | L23R | D | **0.607** | P196Y | D | 0.472 | W82M | D | 0.433 | H143N | N | 0.289 |
| Y18P | D | **0.902** | A28E | D | **0.568** | P197F | D | 0.463 | R23D | D | 0.417 | N297S | N | 0.279 |
| L32H | N | **0.902** | T26A | D | **0.51** | P197W | D | 0.463 | R23G | D | 0.417 | T109S | N | 0.172 |
| L32P | N | **0.902** | G86C | D | **0.508** | R23Q | D | 0.461 | R23L | D | 0.417 | C270A | N | 0.116 |

TS and AspAT both catalyze the transformation of more than one substrate; thus many of the introduced mutations were evaluated to test the effect of enzymatic activity on different substrates. Although data on AspAT were collected for a total of 44 mutations, many of those represent the same substitutions tested over as many as four different substrates, 2-oxoglutarate, L-Asp, L-Glu, and oxalacetate. Similarly in TS, some mutations were characterized with both deoxyuridine monophosphate (dUMP) and tetrahydrofolate (THF) as substrates. **Table 3** distinguishes between the substrates for TS but not for AspAT because dUMP and THF bind to different pockets on TS and therefore require the identification of two different active site coordinates. In contrast, a single binding site is shared among all AspAT substrates; thus only one active site coordinate set is required.

The performance of the prediction procedure can be evaluated from the scoring of enzymes in **Table 3**. The 20 most confident predictions for T4L were called as deleterious (Pred ≥ 0.902). However, only ten of these predictions are supported by experiment, while the remaining ten are not. The outcome is strikingly different for HIV protease where 19 of the 20 mutations scored as deleterious are supported by experimental data, although there is less agreement among the HIV classification trees. A likely explanation for the observed difference in predictability of deleterious mutations between T4L and HIV protease is that only 14.7% of the reported T4L mutations result in the deleterious phenotype compared to 57.1% for HIV protease. Thus there is a much higher probability that a deleterious prediction made for the latter enzyme will be correct. All 14 (eight with dUMP and six with THF) of the top-scoring mutations for TS were called correctly as deleterious. The accuracy of predictions for TS is, as discussed above, due to the large fraction of deleterious mutations out of all substitutions on made on TS^†^. Although only 40.9% of the experimentally characterized AspAT mutations are deleterious, six of the seven predicted deleterious mutations for AspAT were scored correctly, while three substitutions were incorrectly predicted to display a neutral phenotype. The success on AspAT predictability is notable, because for this enzyme, unlike all the others except T4L, the probability that an arbitrarily chosen mutation from the experimental data set is deleterious is much less. This is likely due to the inclusion of the large T4L training set for AspAT predictions.

Of the enzymes examined in this work, only T4L and HIV protease were mutated broadly and without prior consideration of mechanism or distance from the active site. Given the similarity in approach for the T4L and HIV protease experimental studies, the explanation for the much higher frequency of observed deleterious mutations in the latter enzyme remains elusive. Mutations in the other enzymes under consideration here were introduced deliberately, most often with the specific objective of probing the enzyme mechanism. Consequently, most were introduced at or near the active site and preselected as likely to be deleterious.

### Analysis of distance

While the rules for implementation of the BLOSUM and PSAAFS parameters are clearly defined, the contribution of the distance parameter may depend on the accurate localization of the active site thus decreasing its general applicability. In order to explore the sensitivity of this parameter to the precise active site identification, the active site coordinates of HIV protease and T4L, as defined in the Methods, were systematically shifted ±2 Å independently along each of the Cartesian coordinates. These two enzymes were selected for the test because the available data sets provided the most thorough variation on distance. The coordinate variation affected performance by no more than a difference of 0.03 AUC (see **Figure S2**); therefore the contribution of the distance predictor to performance is quite insensitive to the exact placement of the active site.

The precise effect of distance of the mutation from the active site on the predictive accuracy of the method is illustrated by the representation of the T4L mutation data in **Figure 5**, which shows the distance distribution of T4L mutational phenotypes (**see Figure legend for details**). **Figure 5a** explores the distance dependence for all reported mutations. The possibility that active site residues alone dominate the observed trend can be seen from **Figure 5b and c**, which respectively show all data excluding those contacting the bound ligands, N-acetylmuramic acid (MUB) and N-acetyl-D-glucosamine (NAG), and those that are < 8Å from the active site (PDB ID: 148L). The trend line follows a slope that significantly deviates from 0 (P-value < 0.02) in all cases; therefore the distance from the active site is an important predictor of whether a given mutation will seriously degrade catalytic activity. Each bar in **Figure 6** depicts the number of residues at 5Â distance intervals from the active site. The numbers at the top of each bar represent the fraction of deleterious mutations within that interval. That fraction decreases sharply with distance up to 15-20Å, and is relatively constant at about10% beyond that distance.

**Figure 5.**
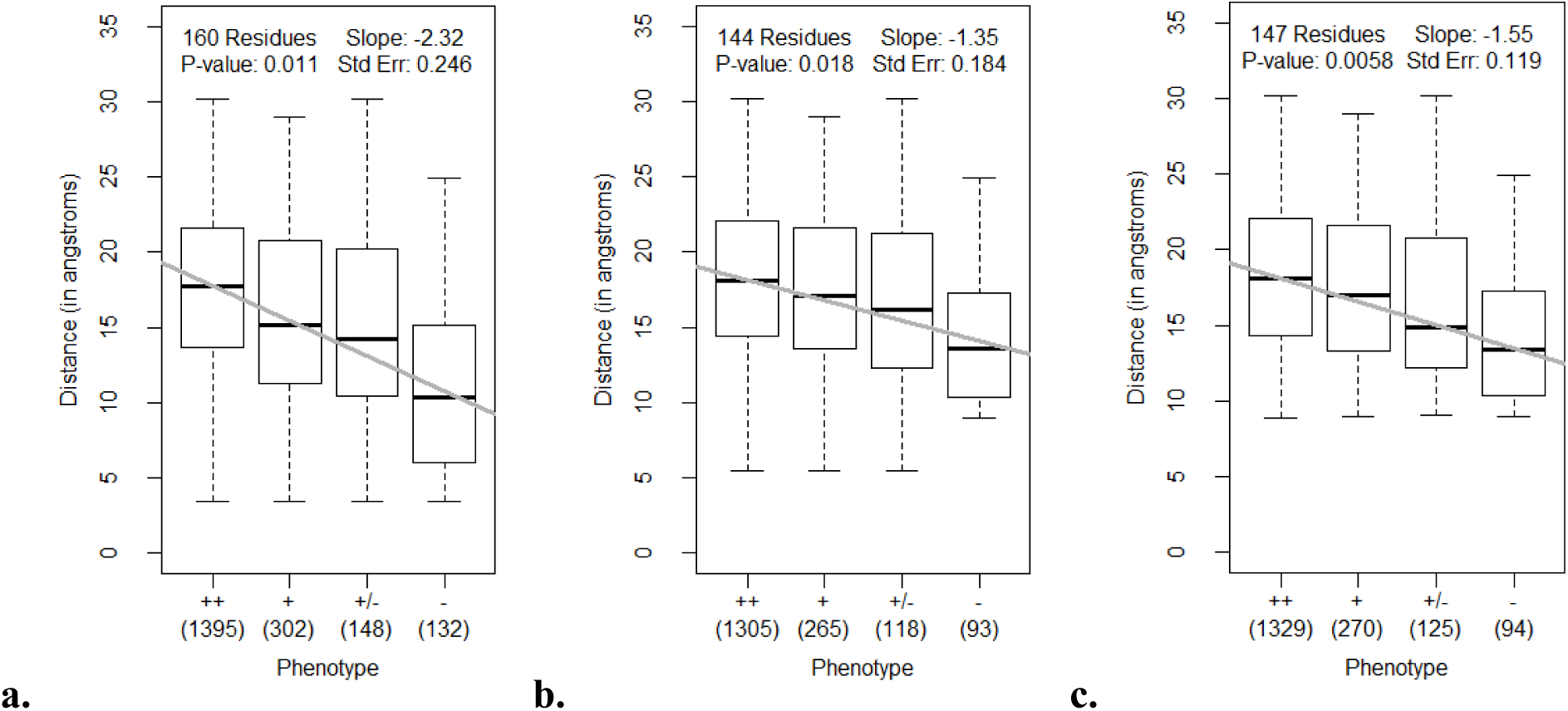
Distribution of distances of T4L mutations from the active site for each observed T4L phenotype. The boxplots organize **(a)** all T4L mutations in the data set, **(b)** omitting residues less than 5Å from the MUB and NAG ligands^68^, and **(c)** excluding residues less than 8Å from the active site (as defined in Methods) into quartiles. The tails denote the minimum and maximum range of each distribution. The bottom and top edges of the box designate the first (25^th^ percentile) and third (75^th^ percentile) quartiles, respectively. The bold division at the center of the box shows the median of the sample. The values below the phenotype labels report the number of mutations within each category. Each gray line represents the weighted linear regression based on the four median values. All P-values are below 0.02, supporting the hypothesis that the slopes ≠ 0. Mutations that introduced (++) and (+) phenotypes are classified as neutral, and those with the (+/-) and (-) phenotypes are classified as deleterious. A total of **(a)** 160 residues with 1977 total mutations, **(b)** 147 residues with 1818 mutations, and **(c)** 1818 mutations covering 147 residues are represented in the plots. Experimental data were collected from Rennell et al^5^.

**Figure 6.**
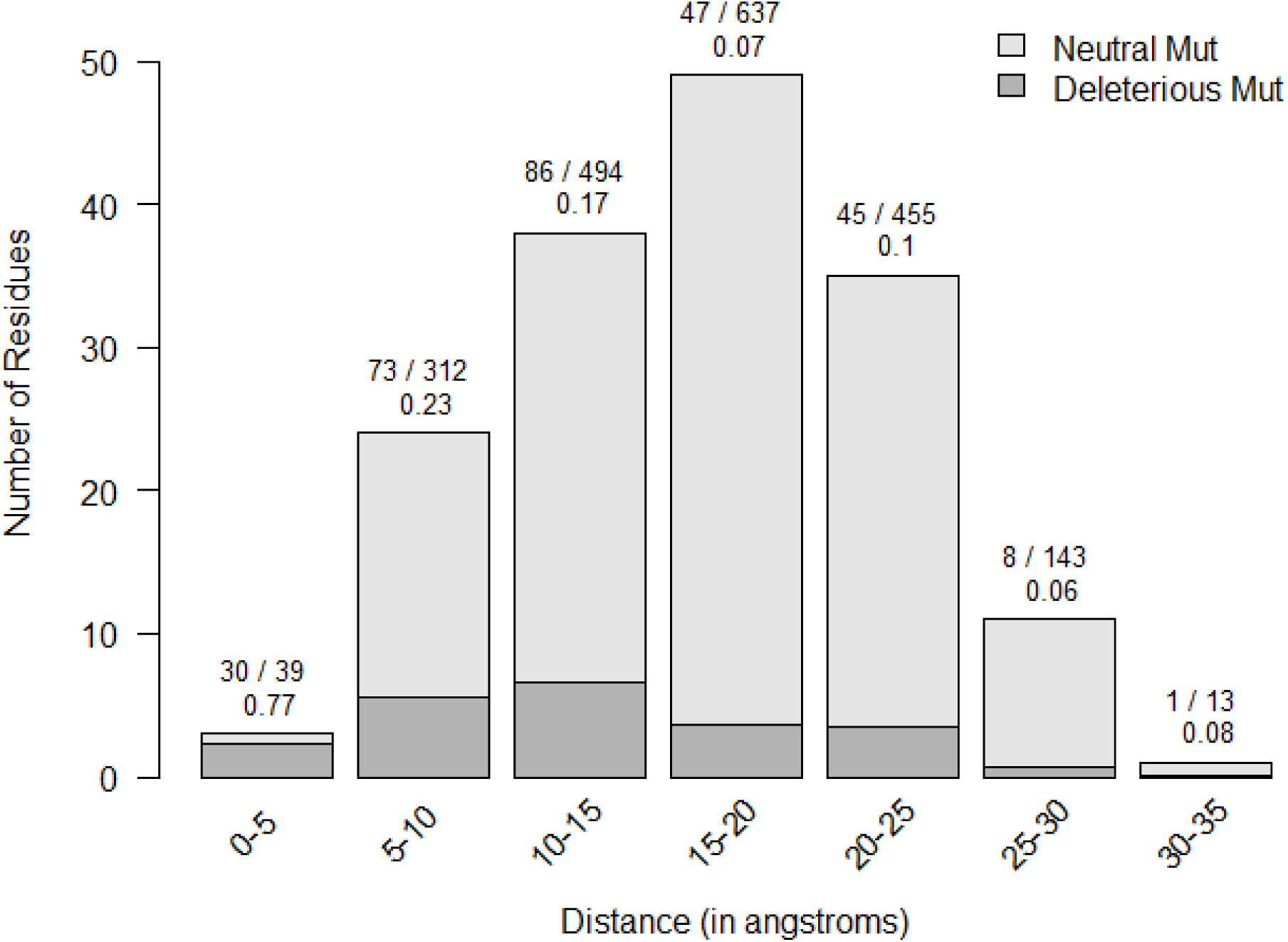
Numbers of residues and of deleterious mutations as a function of the distance from the active site of T4L. In general, every residue in T4L was substituted by 13 different replacements^5^. Mutations to wild-type were excluded from the figure. The number of residues in each distance interval is described by the height of each bar. The dark gray region indicates the total number of deleterious mutations in that range divided by 13, while the light gray section represents the total number of neutral mutations divided by 13. Graphically, this illustrates the ratio of deleterious to neutral mutations found in a particular distance interval from the active site. The proportion of deleterious mutations is given as a fraction out of total mutations characterized in the range and reported in decimal form above each bar.

**Figure 7** presents two sets of ROC curves for LOEO on T4L and HIV protease mutations using the distance predictor only. The difference in ROC curves for the performance of LOEO, using only distance as the predictor, on all T4L mutations versus on only those greater than 5Å from the ligand, MUB and NAG, are negligible (**Figure 7a**). The same test was applied to HIV protease mutations, and we reached a similar conclusion (**Figure 7b**). Together, this indicates that the performance of predictions for mutations away from the catalytic site is equivalent to that of substitutions near the active site.

**Figure 7.**
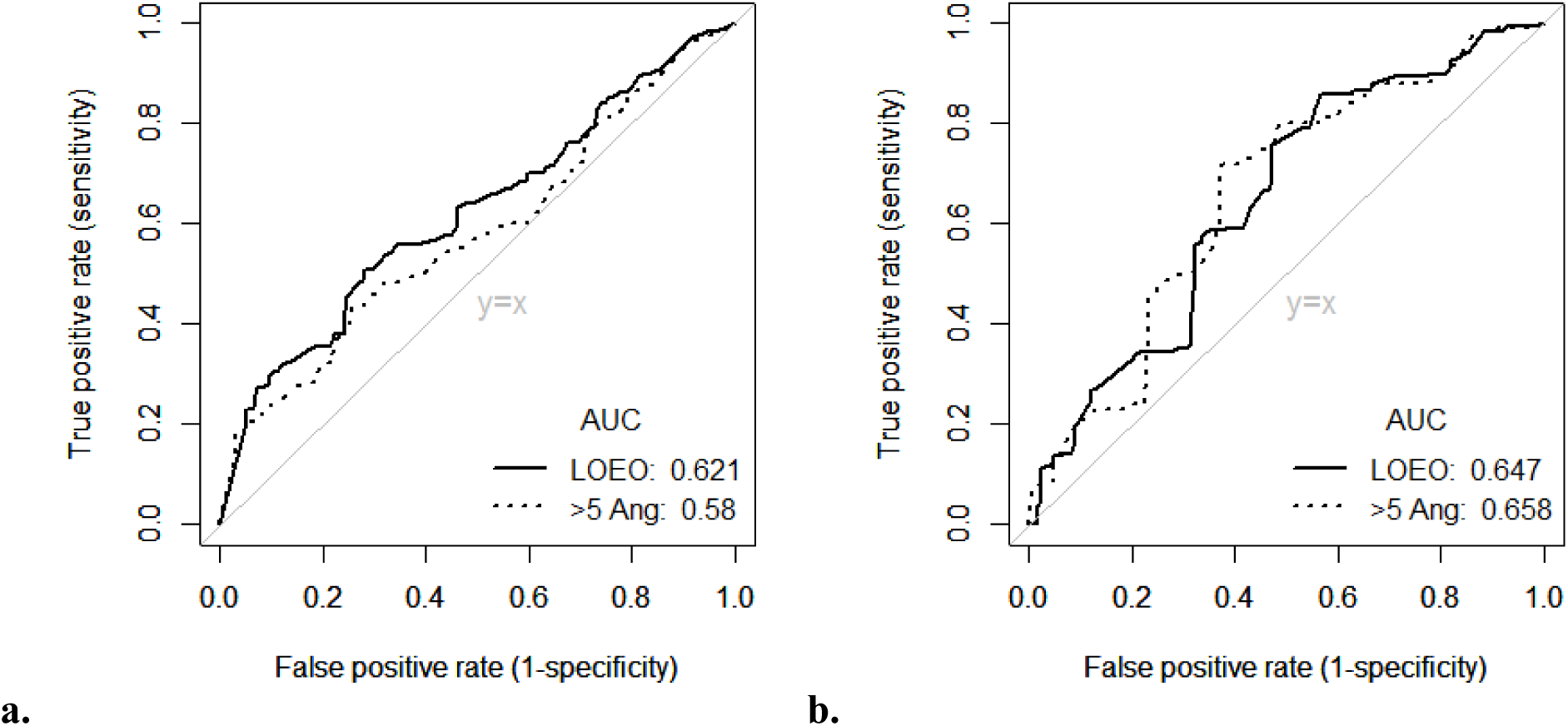
ROC curves for LOEO on T4L and HIV protease mutations using only the distance predictor. T4L and HIV protease mutations are predicted individually using Random Forest trained on mutations on the 11 other enzymes (LOEO). ROC curves illustrating the performance of the predictions for T4L **(a)** and HIV protease **(b)** mutations are shown above. The “Pred All” curves report predictions on all T4L or HIV protease mutations, while the “>5 Ang” curves evaluate predictions on T4L or HIV protease mutations greater than 5Å from its respective ligands. For both T4L and HIV protease, the differences in performance using the distance predictor alone to make predictions for all mutations on a selected enzyme versus those omitting residues less than 5Å from the ligand is negligible.

### Are specific amino acid substitutions in T4L more likely to be deleterious?

**Figure 8** shows the number of substitutions made to each of the 13 target residues in the experimental data on T4L. The total number of replacements to a target amino acid is given at the top of each bar. This number is relatively invariant as expected from the experimental protocol. The shaded section of each bar shows the fraction of deleterious mutations observed when a WT residue is replaced by the one shown on the abscissa. Mutations to C followed K, P, etc. have a higher probability of degrading the catalytic activity than do replacements with eg. G, S, or A. This sensitivity varies over a 10-fold (0.35-0.03) range. Cysteine might have been anticipated as the most damaging substitution as its introduction could lead to such molecular mischief as inter- or intramolecular crosslinking. Proline replacements introduce changes in secondary structure, and might be expected to disrupt catalytic activity where the effects of such perturbations are propagated to the active site. Finally it is noted that replacements with the larger side chain amino acids are clearly more damaging than are substitutions with amino acids bearing smaller side chains. This leads to the conclusion that introduced space is more tolerated on average than is added bulk. These observations, albeit based on a large data set for a single enzyme, do support the use of alanine-scanning mutagenesis as a general probe to evaluate the importance of a given amino acid^52^.

**Figure 8.**
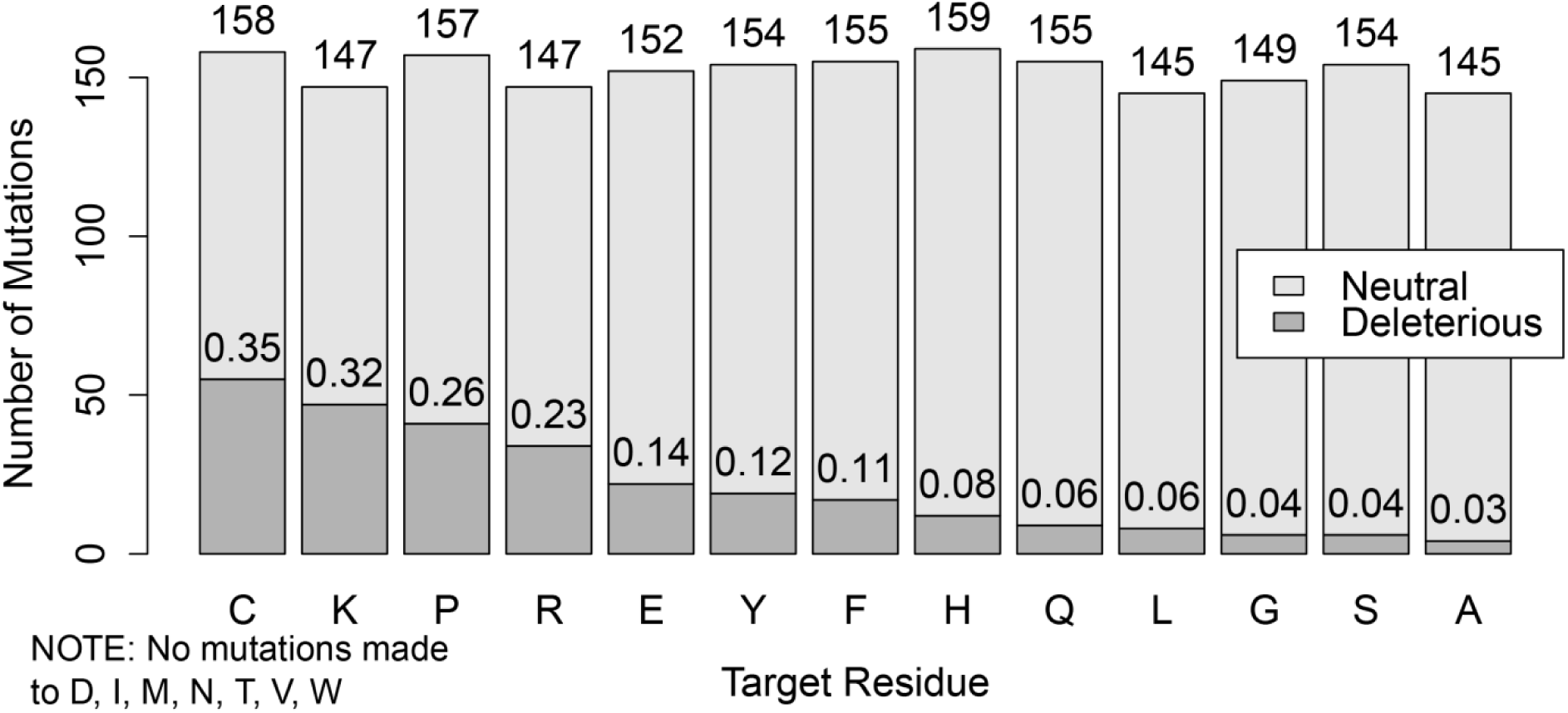
Catalytic effect of targeted mutational changes on T4L, arranged in descending order of the fraction of deleterious mutations made to the specified residue. A total of 1977 mutations were made, 280 of these were deleterious, with at least 97% of the catalytic activity lost upon substitution. The heights of the light gray bars indicate the total number of neutral mutations made to that residue, while the dark gray portions indicate the number of deleterious mutations. Mutations made to C, K, P, or R were the most frequently deleterious (> 20%), whereas mutations made to A, S, or G were rarely deleterious (< 5%). Mutations to alanine were deleterious only ~3% of the time (4 deleterious/145 mutations).

## Footnotes

† The catalytic activity of the H199A mutation was evaluated in two studies separated by 6 years from the Santi laboratory^11,35^). The earlier report was that the mutation decreased the k_cat_/K_m_ value for both substrates by only a factor of ten. This would be scored as neutral in the present classification. The later paper reported that the kinetic parameters were decreased by >500-fold, and is scored as deleterious in accord with our prediction. Dr. Santi (personal communication) wrote that the earlier data were likely compromised by the presence of a small amount of WT contaminant. We relied on his evaluation.

## SUPPLEMENTARY MATERIAL

**Table S1.** List of active site coordinates used to compute the distance predictor.

| Enzyme | Active site coordinate | Details | PDB ID |
| --- | --- | --- | --- |
| T4 lysozyme (T4L) | (10.805, 50.494, 40.451) | A point equidistant to the midpoint of OE1 and OE2 on Glu-11 and the midpoint of OD1 and OD2 on Asp-20 | 148L |
| HIV protease | (15.868, 24.143, 18.502) | Carbonyl carbon (C) between P1 and P1' residues | 4FL8 |
| Thymidylate synthase (TS) | THF: (17.362, 50.167, -49.225); dUMP: (12.608, 48.981, -53.490) | THF: C6 of ligand 10-propargyl-5,8-dideazafolic acid; dUMP: SG of Cys-198 | 1LCA |
| Dihydrofolate reductase | (32.730, 44.617, 12.129) | Midpoint between C7 of ligand methotrexate and C4N of NADP | 1RX3 |
| Aspartate aminotransferase | (-5.695, 56.058, 23.096) | Atom C4A of ligand pyridoxal-5'-phosphate | 1ASM |
| Glutathione transferase | (32.759, 64.359, 63.469) | Geometric center of Ca's for positions 65, 67, 104, 106, 107, 108, 111, 167, 171 on Chain A and positions 104 and 108 on Chain B | 1JLW |
| Salutaridine reductase | (16.070, 62.882, -12.186) | Geometric center of Ca's for positions 180, 236, and 240 on Chain A | 3O26 |
| Hexokinase | (24.754, 1.807, 64.866) | Geometric center of Ca's for positions 151, 168, 169, 204, 205, 231, 256, 289, and 290 on Chain A | 1V4S |
| Arginase | (40.322, 29.108, 47.216) | Geometric center of Ca's for positions 130, 137, 139, 141, 142, 181, 183, 186, 232, and 246 on Chain A of 1RLA | 1RLA (*R. norvegicus*), 2AEB (*H. sapiens*) |
| Fatty acid amide hydrolase | (17.009, -28.840, 26.398) | Geometric center of Ca's for positions 142, 217, and 241 on Chain A | 1MT5 |
| Cytosine deaminase | (-4.494, 31.577, 29.155) | Geometric center of Ca's for positions 33, 51, 62, 64, 91, 94, 114, 152, and 155 on Chain A | 1P6O |
| Fructose-bisphosphate aldolase | (12.729, 4.294, 4.017) | Geometric center of Ca's for positions 35, 61, 109, 110, 226, 227, 264, 265, 267, 286, 288, 289, and 331 on Chain A | 1DOS |

**Figure S1.**
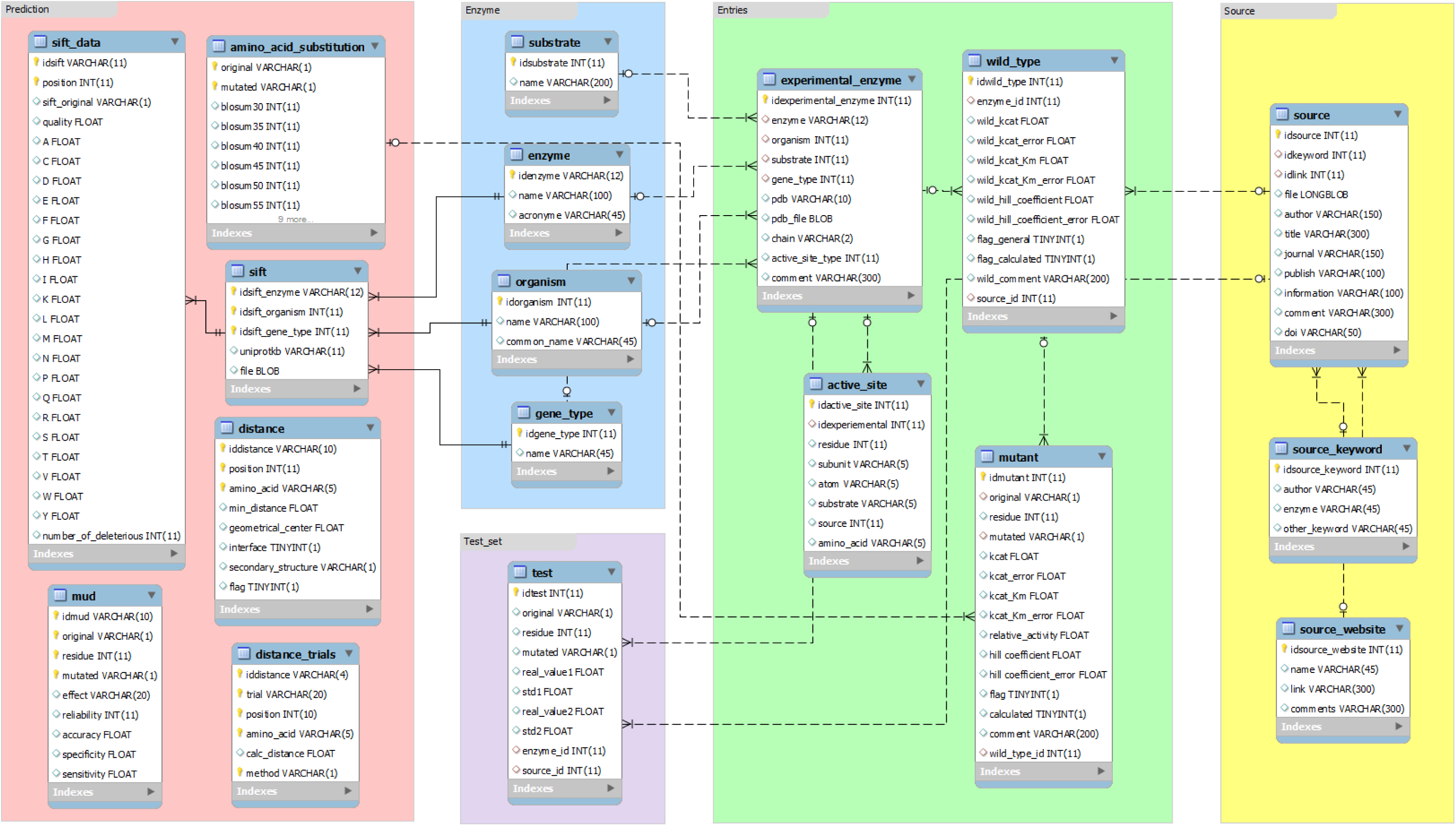
Schema of the mutation data in MySQL. This diagram shows the overall organization of the MySQL database that was used to store mutation-associated data collected from the literature. Each box with a blue border on top is a table with column headers enumerated within the box. The tables are further organized into five main categories: prediction (red), enzyme (blue), entries (green), source (yellow), and test set (violet).

**Figure S2.**
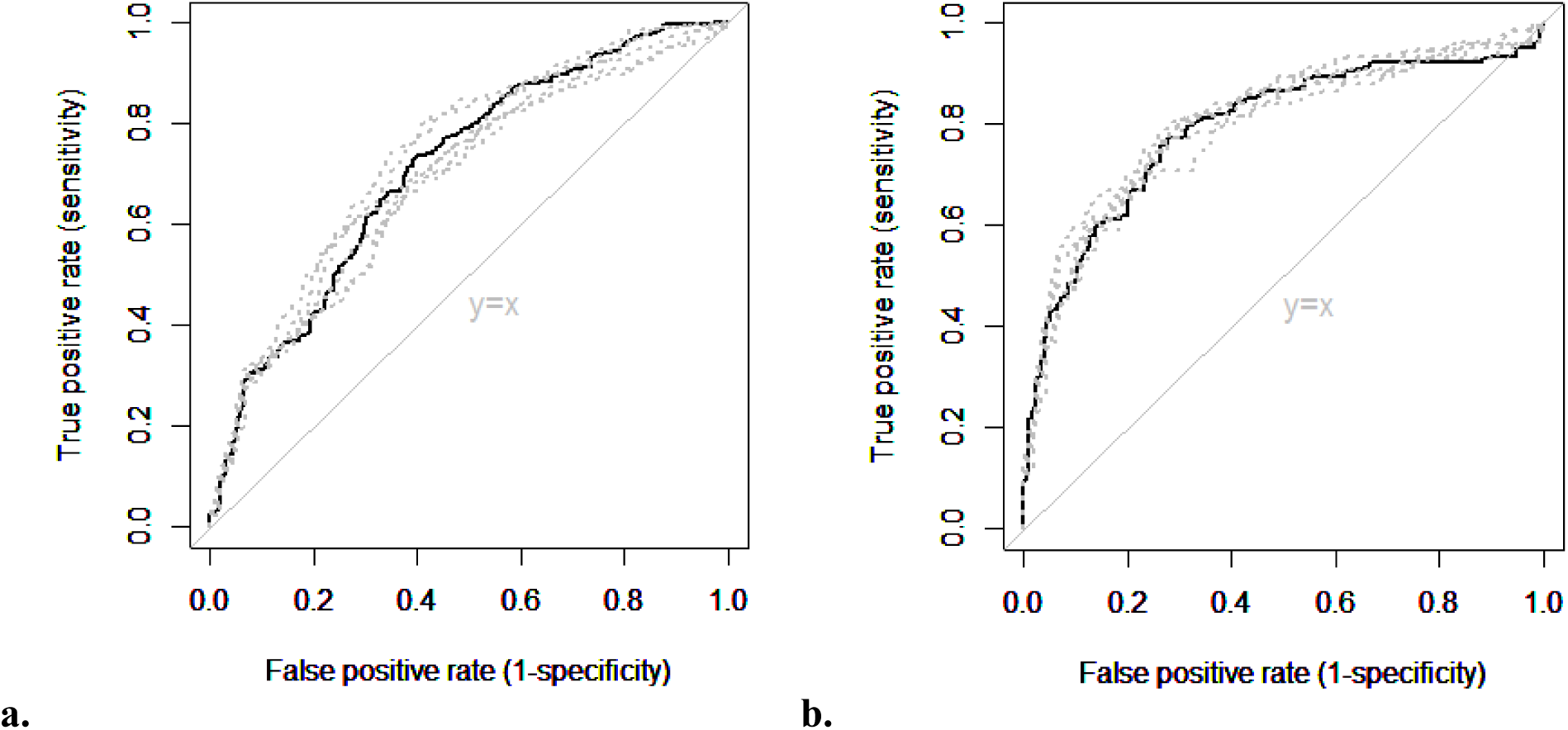
ROC curves measuring the effect of the placement of active site coordinates on method performance. Two sets of ROC curves were constructed, one from the mutation data on **(a)** T4L and the other on **(b)** HIV protease. The black solid curves show the results of 10FCV on the respective enzymes using distance, PSAAFS, and BLOSUM as predictors (0.711 AUC for T4L and 0.791 for HIV protease). The gray dotted curves indicate the results of 10FCV when the active site coordinates are adjusted ±2Å independently along each of the Cartesian coordinates while the other two predictors remain unchanged. The differences between the AUC’s of the solid and dotted curves do not exceed 0.032 for T4L and 0.023 for HIV protease. This demonstrates that the exact placement of the active site coordinate does not affect method performance substantially.

